# Modelling punctuated similarity

**DOI:** 10.1101/2025.07.01.661927

**Authors:** Sarah Kaarina Crockford

## Abstract

Inter-subject, pairwise similarity models provide a methodological resource for flexibly measuring complex, non-linear relationships between brain and behavior. Similarity models, however, can extend beyond brain behavior relationships and can be readily applied to any data where they may be useful. The work presented in this paper introduces a new way of modelling similarity, termed punctuated similarity, where dissimilarity is modelled at a specific point within a given sample of continuous data. With this model researchers can select a value at which they expect dissimilarity or similarity to occur and evaluate it against real similarity distributions. To demonstrate how this model works two separate proof-of-concept experimental analyses were conducted on publicly available data, the results of which are reported in this paper. The first investigated puberty as a critical time-point for growth in a sample of children aged five to nineteen obtained from the NCD Risk Factor Collaboration. The second investigated the COVID-19 pandemic as a critical time-point for market volatility in stock price data for market indices from seven different countries. Both experiments showed that the similarity model presented in this paper was effective at modelling critical time points for increased variance in stock market prices and growth development of children. Finally, this paper is accompanied by an open-source R library to recreate the similarity models presented, providing a tool for future researchers to use in their own analysis.

## Introduction

Heterogenous data, such as data derived from large, diverse cohorts of the human population or clinical subjects, is characterized by a high degree of internal variance and complexity. Therefore, complex heterogenous data relationships between dependent and independent variables may not always exhibit a linear, one-to-one mapping, trajectory. The increasing availability of such large-scale, naturalistic data thus requires quantitative methodological tools that are sensitive to its complexity and diversity (Boylan et al., 2025; Cirillo & Valencia, 2019; Dolley, 2018; Prosperi et al., 2018), such as analytical models that focus on representing the underlying structure of the data to understand where in the data complexity and variance may occur. To this end, similarity-based modeling is a flexible analytical approach that can reveal meaningful relationships across conditions, individuals, or modalities.

Similarity-based models such as MVPA (Haxby et al., 2001) and RSA (Kriegeskorte et al., 2008) have been successfully applied in neuroimaging studies to identify patterns of brain activity in response to visual stimuli (Kriegeskorte et al., 2008; Haxby et al., 2001; Haxby, 2012) and to model salient events in naturalistic conditions, such as during passive story listening or movie watching (i.e., Baldassano et al., 2017; Camacho et al., 2023; Chang et al., 2022; Finn et al., 2020; Nastase et al., 2019). Finn et al., (2020) previously used similarity between subjects on behavioral measures, such as personality traits or working memory, to investigate neural response similarity while subjects passively watch a movie. The authors constructed hypothetical distributions of pairwise behavioral similarity, arranged by subjects’ scores on these behavioral measures, and compared them to the real similarity distributions of neural responses from the same pairs of subjects. For example, if subjects who score very similarly on a working memory task are also similar in patterns of neural activation while watching a movie, then a behavioral model that models subjects who are closer in their scores as also most similar will share a strong relationship with the real neural similarity. These behavioral models can be flexible to whichever distributions a researcher wishes to model. So far, previous work has demonstrated their effectiveness at modelling subjects who are closest in behavior as being most similar, subjects as they increase on a given behavior becoming more similar and, vice-versa, subjects as they increase on a given behavior becoming less similar (Camacho et al., 2023; Finn et al., 2020) (Figure 1).

**Figure 1:**
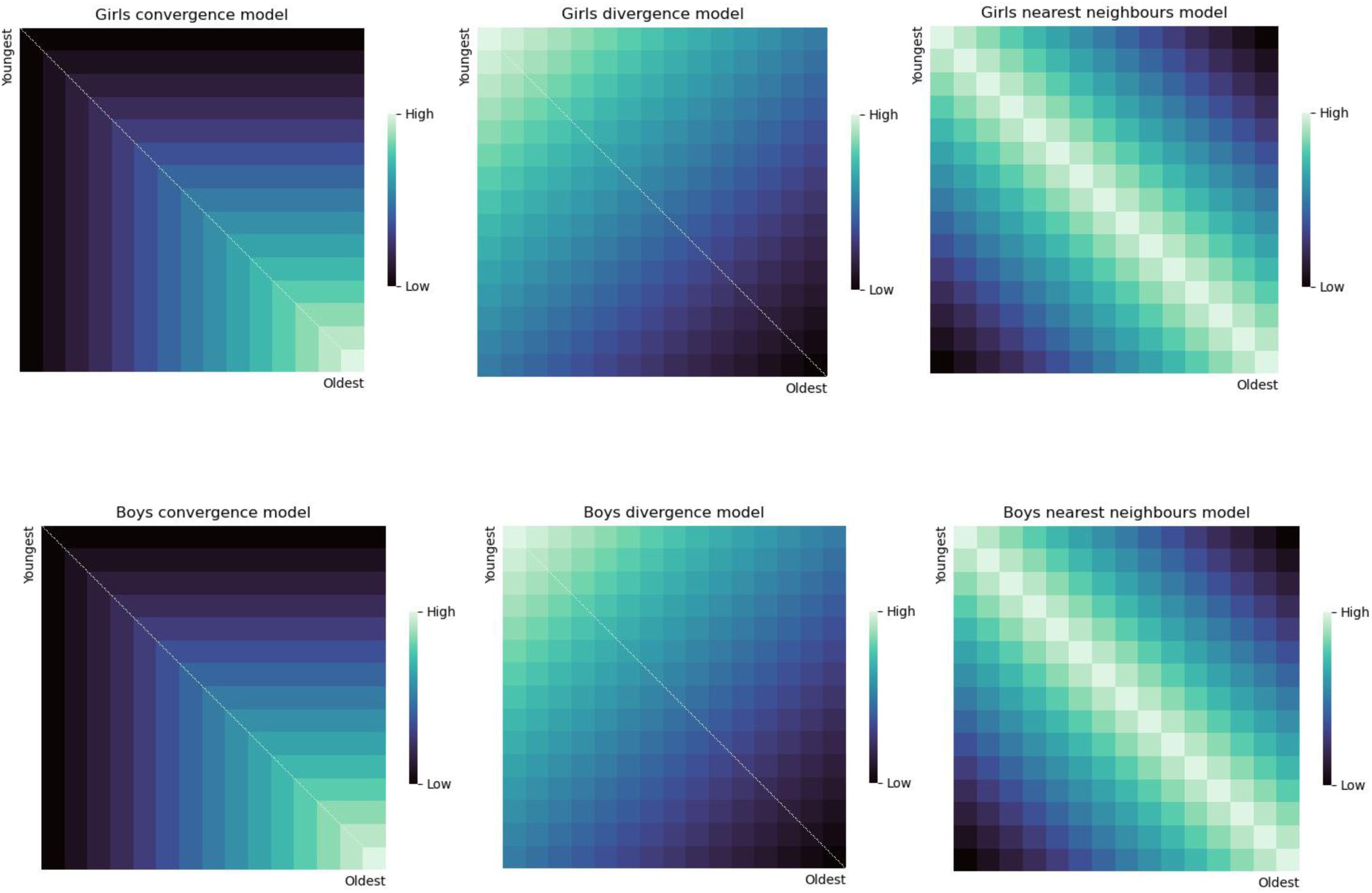
Convergence, divergence and nearest neighbor similarity distributions in girls and boys from the height data used in experiment one of this paper. These models were obtained from the age data provided, where the rows and columns of these matrices are arranged from youngest (5 years of age) to oldest (19 years of age).

This work presented in this paper extends previous similarity work introduced by Finn et al., (2020) and introduces a novel method to model ‘moments’ of dissimilarity or similarity, termed ‘punctuated similarity’. The model owes its name to the notion of punctuated equilibrium, where trajectories such as evolutionary change are marked by bursts of activity preceded and followed by steady changes (Eldredge and Gould, 1972; Yackinous, 2015). The punctuated similarity model requires that a researcher has a prior hypothesis of where they might expect a moment of dissimilarity or similarity to occur and then apply it to the model, meaning it uses a confirmatory approach to establish where moments of heightened dissimilarity might occur. To evaluate the proposed punctuated similarity models, data is presented from two experiments. In the first experiment, these models are examined in relation to global height growth data in children aged five to nineteen, where it is widely known that the onset of puberty is a critical moment for rapid changes in height (i.e., Suutela et al., 2022). In the second experiment the models were evaluated against similarity in stock price data from different market indices from seven different countries (United Kingdom, United States, Germany, Italy, France, Australia, and Hong Kong). Specifically, this paper looked at stock price similarity sixth months prior and after the onset of the COVID-19 pandemic, which presented a critical time-point for increased stock market volatility (Kayani et al., 2024). Finally, results from the punctuated model proposed in this paper were formally compared to the other three models (nearest neighbor, convergence, and divergence) proposed by Camacho et al., (2023) and Finn et al., (2020). The methods and results for each experiment are detailed separately below.

### Modelling similarity

The first three similarity models used in this work were replications of Finn et al., (2020)’s and Camacho et al’s (2023) work (for their code, see here: https://github.com/esfinn/intersubj_rsa; https://github.com/catcamacho/hbn_analysis), namely ‘nearest neighbor’, ‘convergence’, and ‘divergence’. Nearest neighbor was modeled by taking the maximum sample value for age and, from it, subtracting the absolute difference in age between each subject pair. Convergence was modelled as the minimum value between the two age values for a pair of subjects and divergence was modelled as the maximum sample value for age, minus the average age in each pair of subjects.

Results of the previous approaches were compared to the new model proposed in this paper (termed ‘punctuated similarity’), which allows one to model dissimilarity where the tail ends of an extreme (i.e. younger and older subjects) might be more similar, compared to subjects who are centered around a given value (i.e., children around the age of ten). The equation used to model this similarity distribution is presented below:

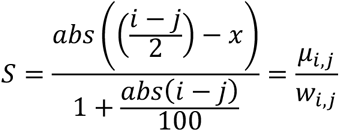

Where S is the final similarity value, i is the given value for subject i and j is the given value for subject j, x is the value around which we want to center our similarity distribution and w is the weighted difference. In sum, punctuated similarity is computed using the absolute difference between the mean age (value) for each pair of subjects and a ‘center’ value, that is a value around which similarity or dissimilarity should be centered. For example, to model that the most dissimilar subjects are those closest to the age of 10, with increasing between subject similarity as age either increases or decreases from 10, the center value is then 10 years of age. Weighting this value by the absolute difference between the pair values (i, j) ensures that similarity remains in the upper left and bottom right corners of the matrix (Figure 2).

**Figure 2:**
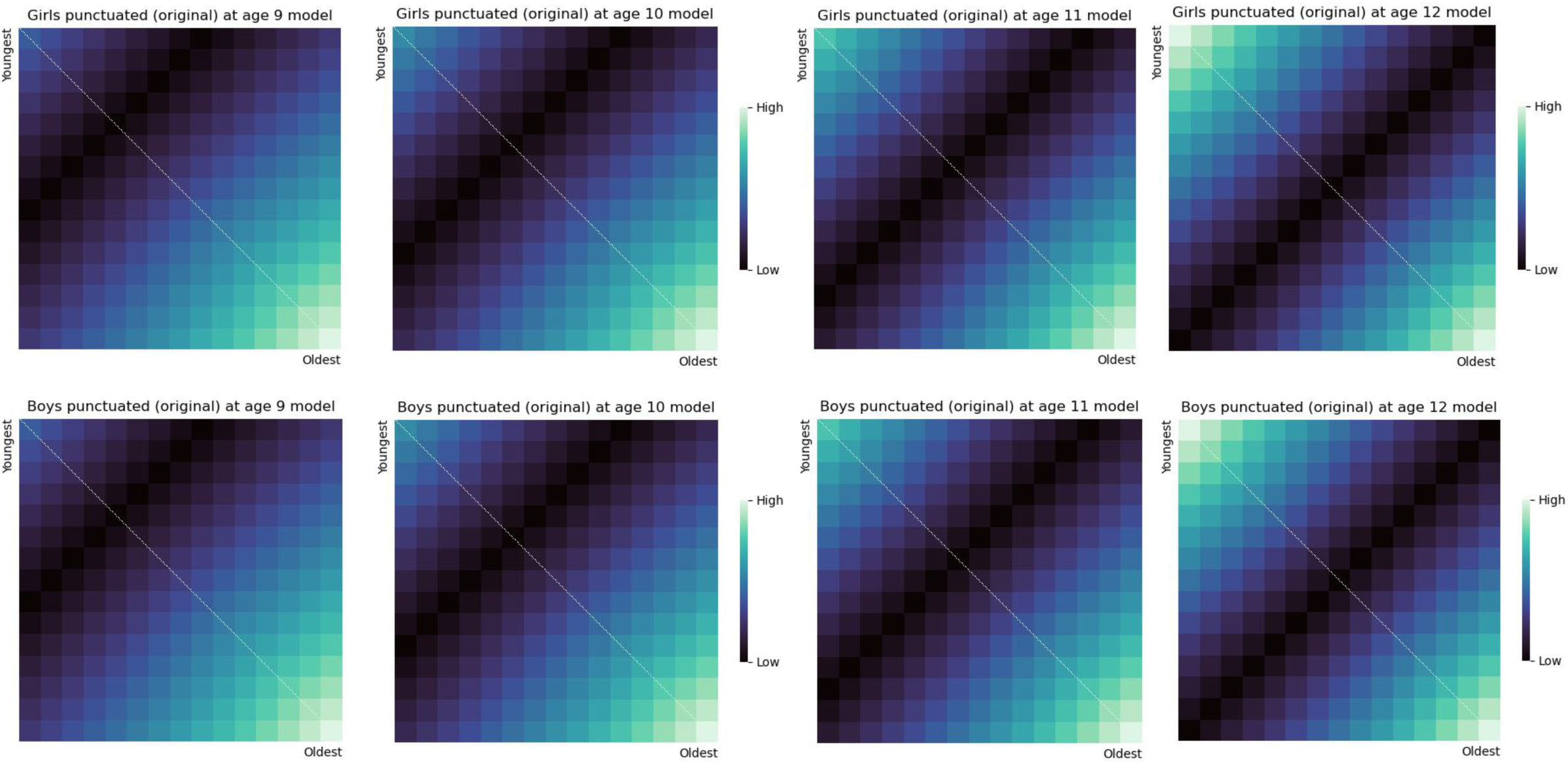
Punctuated similarity distributions in girls and boys from the data used in this paper, modelled at the ages of 9, 10, 11 and 12 (from left to right). These models were obtained from the age data provided, where the rows and columns of these matrices are arranged from youngest (5 years of age) to oldest (19 years of age) child.

In this conceptualization of similarity, dissimilarity becomes very strong at the value around which it is centered. However, it may be the case that even though that center value represents a moment of dissimilarity, there is still similarity between subjects closest on that metric. Which is to say, that there is still subtle evidence of a nearest neighbor effect, although attenuated by dissimilarity around a centered value. Therefore, this paper also presents a further fifth model that extends the notion of punctuated similarity but also balances out a potential nearest neighbor effect. To do so, one simply sums the punctuated similarity computation to the nearest neighbor computation (Figure 3).

**Figure 3:**
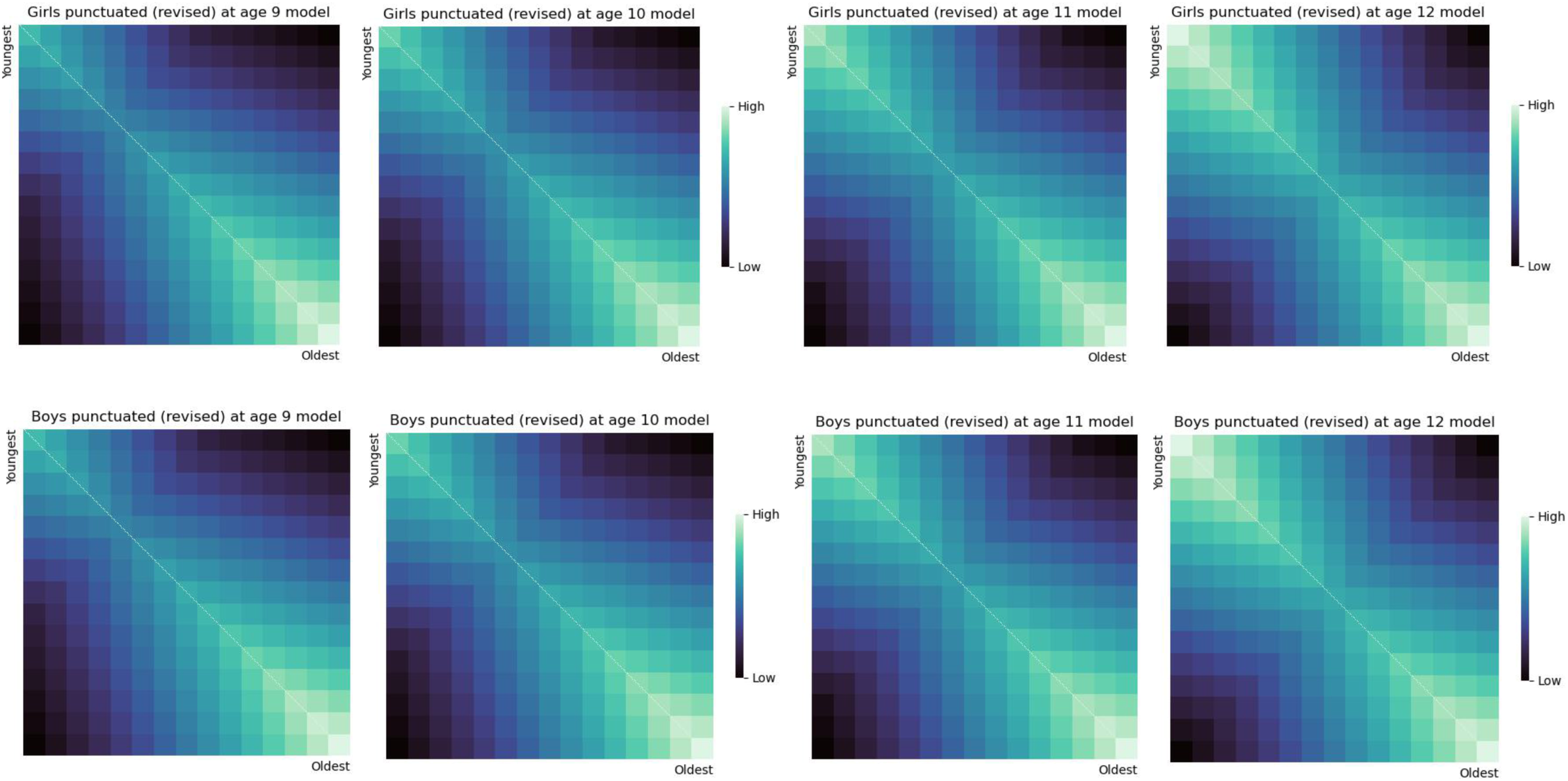
Revised punctuated similarity distributions which include the nearest neighbor effect in girls and boys from the data used in this paper, modelled at the ages of 9, 10, 11 and 12 (from left to right). These models were obtained from the age data provided, where the rows and columns of these matrices are arranged from youngest (5 years of age) to oldest (19 years of age) child.

### R Library ‘similarity models’

For future usability and reproducibility purposes, this paper is accompanied by an openly available R library containing the functions necessary to build the above similarity matrices on any given one-dimensional set of numeric/integer data (https://github.com/sarahkaarina/similaritymodels). The R library includes the nearest neighbor and convergence models developed by Finn et al., (2020), which are released under the MIT license. Credit to the authors is provided in the documentation for the ‘similaritymodels’ library.

## Experiment 1: Height and the onset of puberty

The onset of puberty marks a critical time point in childhood related to rapid changes in several biological mechanisms including but not limited to height, metabolism, neural structure and functioning, and language development (Karlberg et al., 2023; Suutela et al., 2022; Rosselli et al., 2014; Tanner et al., 1976; Taranger & Hägg, 1980; Vijayakumar et al., 2018). Importantly, the onset of puberty is often defined by the age at which changes in height reach their peak velocity; that is, the age at which children begin their rapid growth spurt (Suutela et al., 2022). The onset of puberty can range from nine to fifteen years of age (Dorn et al., 2006; Vijayakumar et al., 2018), with average estimates for the peak in growth for girls being earlier (about 11 years of age) than boys (about 13 years of age) (Suutela et al., 2022). Indeed, height is arguably the “most significant *nonlinear* feature in the growth process” (Tang et al., 2024 on pg. 1, quoting Tanner, 1978). This would suggest that the earliest onset of puberty should relate to increased dissimilarity in heights, as children arrive at and reach their peak growth at different time-points.

Therefore, if the model proposed in this paper is effective at measuring dissimilarity in heights during puberty, then, compared to other models proposed by Finn et al., (2020) and Camacho et al., (2023), it should share the strongest relationship with the real similarity distribution of heights. Additionally, if the punctuated similarity model is sensitive to differences in timings between boys and girls, then a model centered around an earlier age should be most significant for girls compared to a model centered around a later age for boys. To test this hypothesis, open-source average height data for boys and girls aged 5 to 19 obtained from the NCD Risk Factor Collaboration (2023) was used to compute height-based similarity matrices, arranged from youngest to oldest child. Using previous methods for comparing similarity matrices (Camacho et al., 2023; Finn et al., 2020; Mantel, 1967), modelled similarity was then related to the real similarity matrices. To assess which age of pubertal onset best modeled height related dissimilarity, the punctuated model was centered starting from the age of nine (earliest age of onset, Dorn et al., 2006) and stopping at the age twelve (the age just before boys reach peak growth, Suutela et al., 2022).

## Experiment 1: Methods

### Data

Data for this paper was downloaded from the NCD-RisC repository (https://ncdrisc.org/data-downloads-height-urban-rural-ado.html; last accessed: 15/06/2025) (NCD Risk Factor Collaboration, 2023). Mean height data obtained from 71-million children aged 5 to 19 sampled from urban and rural areas in 200 countries between the years 1990 to 2020 was used (see NCD Risk Factor Collaboration, 2023 for a full description of the data and related collection/sampling procedures). For each year sampled (N = 31, from 1990 to 2020) and for each age (N = 15, from 5 to 19), the data file obtained from NCD-RisC repository had a related mean height, recorded in centimeters. Data was then categorized by belonging to male (boys) or female (girls) samples and whether they were sampled from rural or urban populations. This resulted in a total of 465 mean height values for each data category (urban boys, rural boys, urban girls, or rural girls), where each year contained fifteen mean height samples (31 years x 15 age values = 465) (Figure 4).

**Figure 4:**
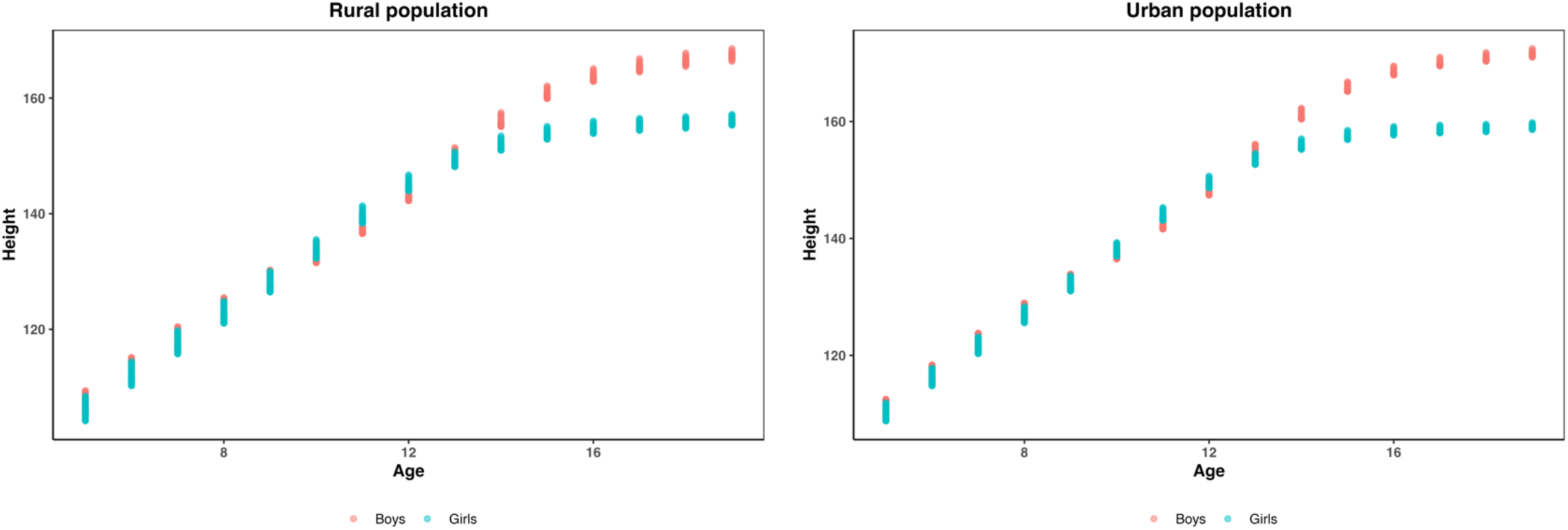
Age (years) in relation to height (centimeters) for rural (left figure) and urban (right figure) populations, where blue dots represent girls and red dots represent boys.

### Computing height related similarity

Similarity between height was computed as the absolute difference between each given height value within a category (urban boys, rural boys, urban girls, or rural girls), resulting in a diagonal of 0 values (0 being absolute similarity). To maintain consistency with the modelled similarity matrices, the subject-by-subject similarity matrix was rescaled to values between 0 and 1 and these values were then flipped by subtracting them from 1, so that highest similarity (the diagonal) was represented by 1 and 0 represent the most dissimilarity. This computation resulted in four symmetric matrices, one for each category, of 465 rows and 465 columns, arranged by increasing age values so that heights related to younger ages were in the upper left corner and heights associated with older ages were in the bottom right corner (Figure 5).

**Figure 5:**
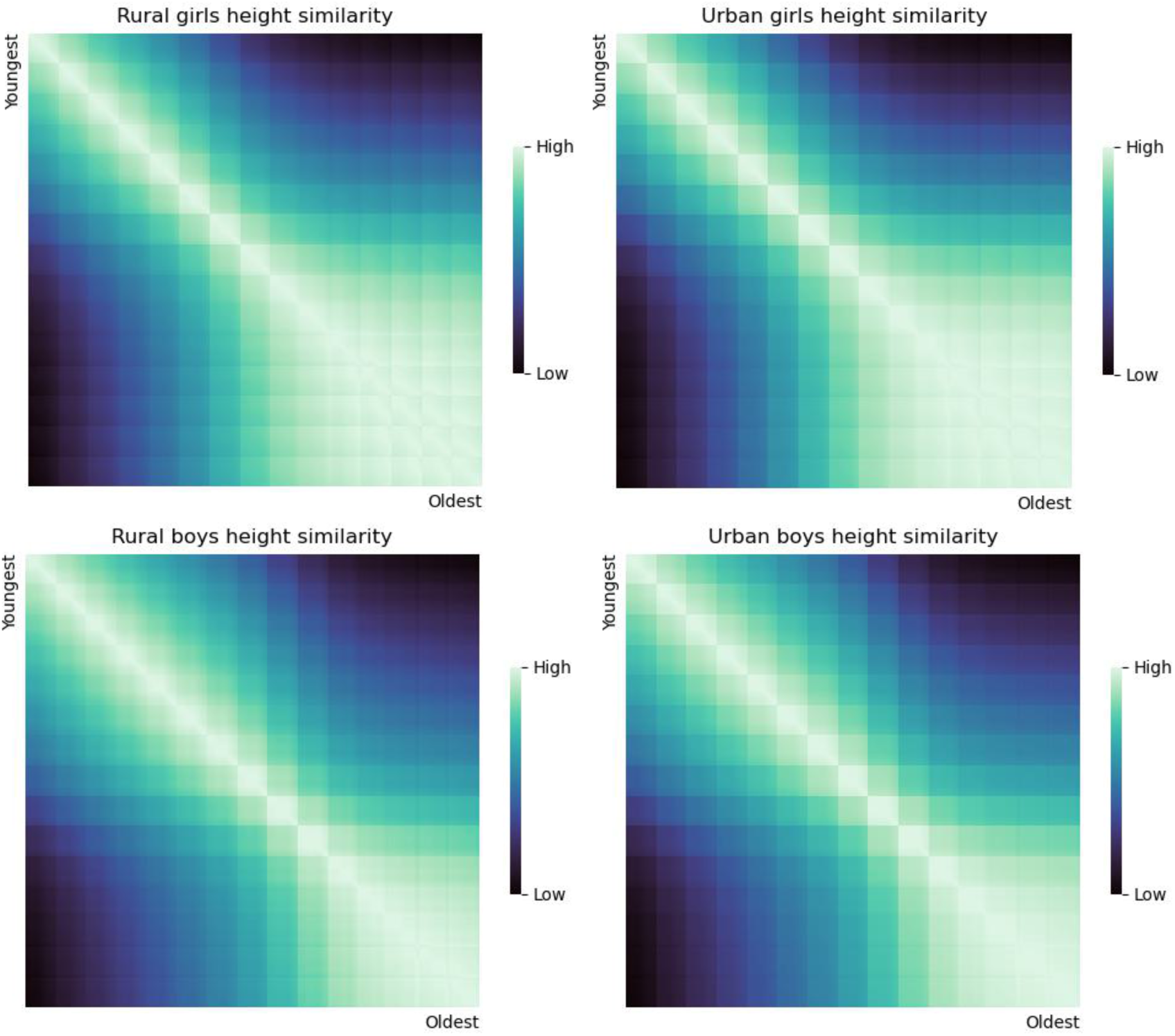
Real similarity distributions for height in urban and rural girls (top) and urban and rural boys (bottom). These models were obtained from the mean height data provided, where the rows and columns of these matrices are arranged from youngest (5 years of age) to oldest (19 years of age) child.

### Relating similarity matrices

To compare modelled similarity with real similarity distributions, a non-parametric mantel test was used (Camacho et al., 2023; Finn et al., 2020; Mantel, 1967) implemented using the ‘vegan’ packaged in R (https://cran.r-project.org/web/packages/vegan/index.html; last accessed: 29/04/2025). A non-parametric mantel test computes a Spearman’s rank correlation between the lower triangles of two similarity matrices and assumes that the matrices are of symmetric and of the same row x column dimension. The mantel test also permuted the rows and columns to generate a random distribution of correlation values and from that obtained a p-value representing how significantly related two matrices are. For replicability and computation speed, 1000 permutations were used (from start to finish, the analysis from each experiment reported in this paper takes approximately 30 to 45 minutes to complete).

To determine which comparisons were significantly different from each other (p-value < 0.05), each resulting Spearman’s Rho from the mantel test was compared using a Fisher’s z comparison of correlation coefficients (Fisher, 1928). For boys and girls, this analysis collapsed the urban and rural groups as to formally compare not only which similarity model best related to the real height similarity data, but also to check whether this was consistent for both urban and rural populations. The motivation behind this was to use the urban and rural data as ‘discovery’ and ‘replication’ datasets to confirm the results from the modelled similarity correlations. Although differences in growth between urban and rural populations are reported (NCD Risk Factor Collaboration, 2023; Tang et al., 2024), the punctuated similarity should still be the most related to height similarity compared to other modelled similarity distributions, even if at different onset ages for puberty.

## Experiment 1: Results

### Punctuated similarity modelled around the age of 11 best relates to height similarity in boys and at the age of 9 for girls

As evident in Tables 1 and 2, nearly every model, excluding divergence, shared a significant relationship with the real height similarity distributions across all groups (urban boys, rural boys, urban girls, or rural girls). To understand which models shared the strongest relationships, this analysis used a Fisher’s test to compare each resulting R value for each comparison in the boy and the girl groups separately. The p-values from the resulting 231 comparisons (eleven age by height comparisons for two groups, urban and rural) were adjusted using the Benjamini-Hochberg False Discovery Rate (BH-FDR) correction method for boys and girls separately. After ordering the data by highest R values, only the highest R values that were not statistically significant to each other but statistically significant to every other comparison were considered. That is, all the R values that were significantly higher than every other R value but were not statistically significant different from other high R values were considered ‘winners’ in this analysis (Figure 6).

**Figure 6:**
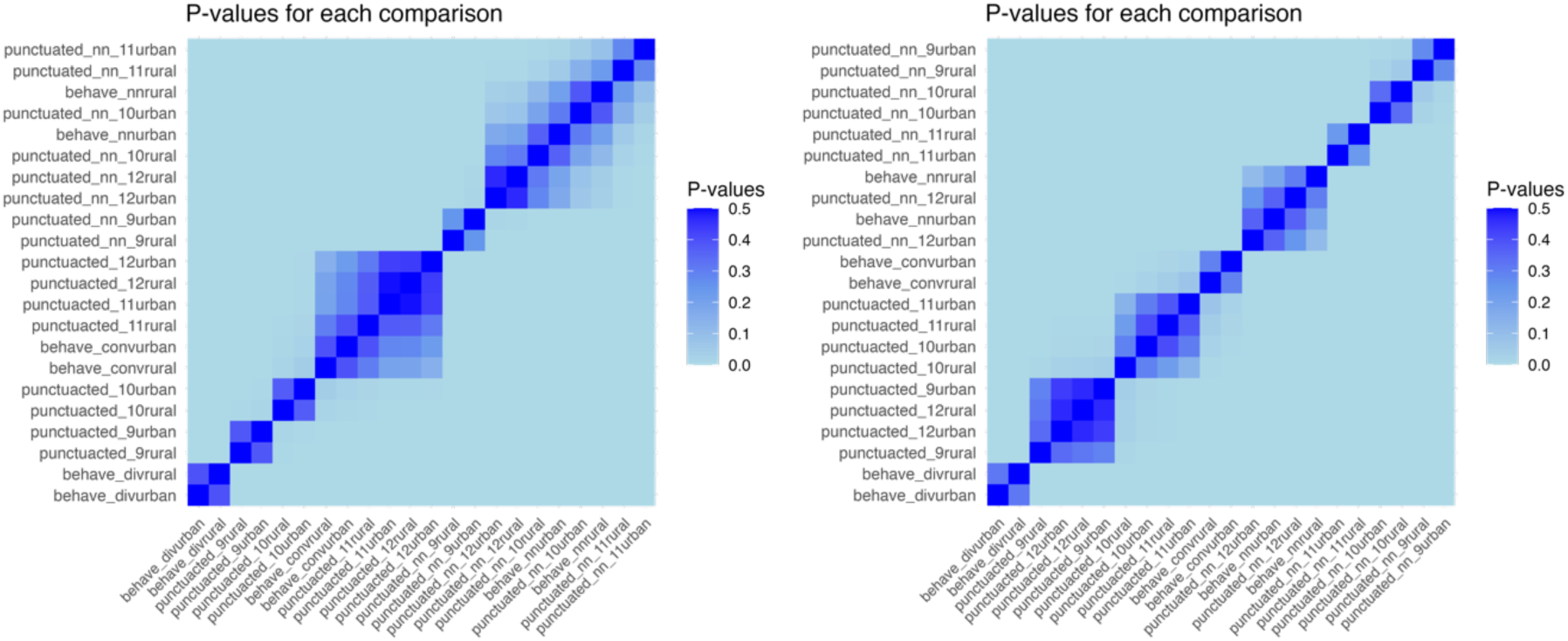
Matrix of resulting p-values from each R comparison for boys (left) and girls (right), arranged by increasing R values along the column (highest R values are in the upper left and bottom right corners). The highest R values were considered those that ‘clustered’ in the upper right corner, where they were significantly different from all the other R values but had no remaining higher R values that significantly differed from them. As it is possible to see for the boys (left figure), this upper right corner cluster contained the resulting R values for our revised punctuated similarity model centered at age 11 and the nearest neighbor model for the rural boy population. For the girls (right figure), this upper ight corner cluster only featured the revised punctuated similarity model centered at age 9.

**Table 1:** Results from the boys’ group. *R values bolded in red were significantly higher than all other reported R values.*

|  | Urban R | Urban Adj. P-value | Rural R | Rural Adj. P-value |
| --- | --- | --- | --- | --- |
| **Nearest Neighbor** | 0.959 | 0.001 | **0.963** | **0.001** |
| Convergence | 0.604 | 0.001 | 0.592 | 0.001 |
| Divergence | -0.190 | 1.0 | -0.172 | 1.0 |
| Punctuated (age 9) | 0.362 | 0.001 | 0.343 | 0.001 |
| Punctuated (age 10) | 0.508 | 0.001 | 0.490 | 0.001 |
| Punctuated (age 11) | 0.630 | 0.001 | 0.616 | 0.001 |
| Punctuated (age 12) | 0.638 | 0.001 | 0.631 | 0.001 |
| Punctuated revised (age 9) | 0.936 | 0.001 | 0.930 | 0.001 |
| Punctuated revised (age 10) | 0.962 | 0.001 | 0.956 | 0.001 |
| **Punctuated revised (age 11)** | **0.970** | **0.001** | **0.967** | **0.001** |
| Punctuated revised (age 12) | 0.953 | 0.001 | 0.953 | 0.001 |

**Table 2:** Results from the girls’ group. *R values bolded in red were significantly higher than all other reported R values.*

|  | Urban R | Urban Adj. P-value | Rural R | Rural Adj. P-value |
| --- | --- | --- | --- | --- |
| Nearest Neighbor | 0.868 | 0.001 | 0.882 | 0.001 |
| Convergence | 0.766 | 0.001 | 0.748 | 0.001 |
| Divergence | -0.425 | 1.0 | -0.397 | 1.0 |
| Punctuated (age 9) | 0.587 | 0.001 | 0.560 | 0.001 |
| Punctuated (age 10) | 0.682 | 0.001 | 0.661 | 0.001 |
| Punctuated (age 11) | 0.701 | 0.001 | 0.690 | 0.001 |
| Punctuated (age 12) | 0.579 | 0.001 | 0.583 | 0.001 |
| **Punctuated revised (age 9)** | **0.978** | **0.001** | **0.976** | **0.001** |
| Punctuated revised (age 10) | 0.968 | 0.001 | 0.970 | 0.001 |
| Punctuated revised (age 11) | 0.930 | 0.001 | 0.936 | 0.001 |
| Punctuated revised (age 12) | 0.861 | 0.001 | 0.873 | 0.001 |

Results in the boys’ group determined that the revised punctuated similarity model attenuated for nearest neighbor effects modelled as dissimilarity around the age of 11 resulted in the highest correlation with real similarity in height. This result was consistent in both the urban and rural groups of boys, although in the rural group, the nearest neighbor similarity model also related to the real height data significant more than other models (Table 1). Instead, for girls the revised punctuated similarity model modelled as dissimilarity around the age of 9 shared the highest relationship with real height similarity (Table 2).

## Experiment 2: Market volatility

To further examine the applicability of the proposed punctuated similarity models, these were also applied to stock price data. More specifically, dissimilarity was modelled at the onset of the COVID-19 pandemic related to increased dissimilarity in daily stock prices for a subset of stock market indices from different countries (United Kingdom/FTSE-100, United States/S&P 500, Germany/DAX, France/CAC 40, Italy/FTSE MIB, Hong Kong/HIS and Australia/S&P/ASX 200). Previous research has already demonstrated that the onset of the COVID-19 pandemic was a critical moment for heightened stock volatility worldwide (Kayani et al., 2024). Therefore, stock prices around March 2020, the onset of the pandemic (Spiteri et al., 2020), should change rapidly from day to day, compared to moments, such as prior and after the initial stages of the pandemic, where the daily change in stock price is not so steep. To demonstrate this, the R library ‘quantmod’ was used (https://github.com/joshuaulrich/quantmod; last accessed: 17/06/2025) to plot the daily stock prices at open and closing times for the seven market indices used in this analysis, where a clear drop off in stock price followed by a return in stability is evident around the March 2020 timepoint (Figure 7).

**Figure 7:**
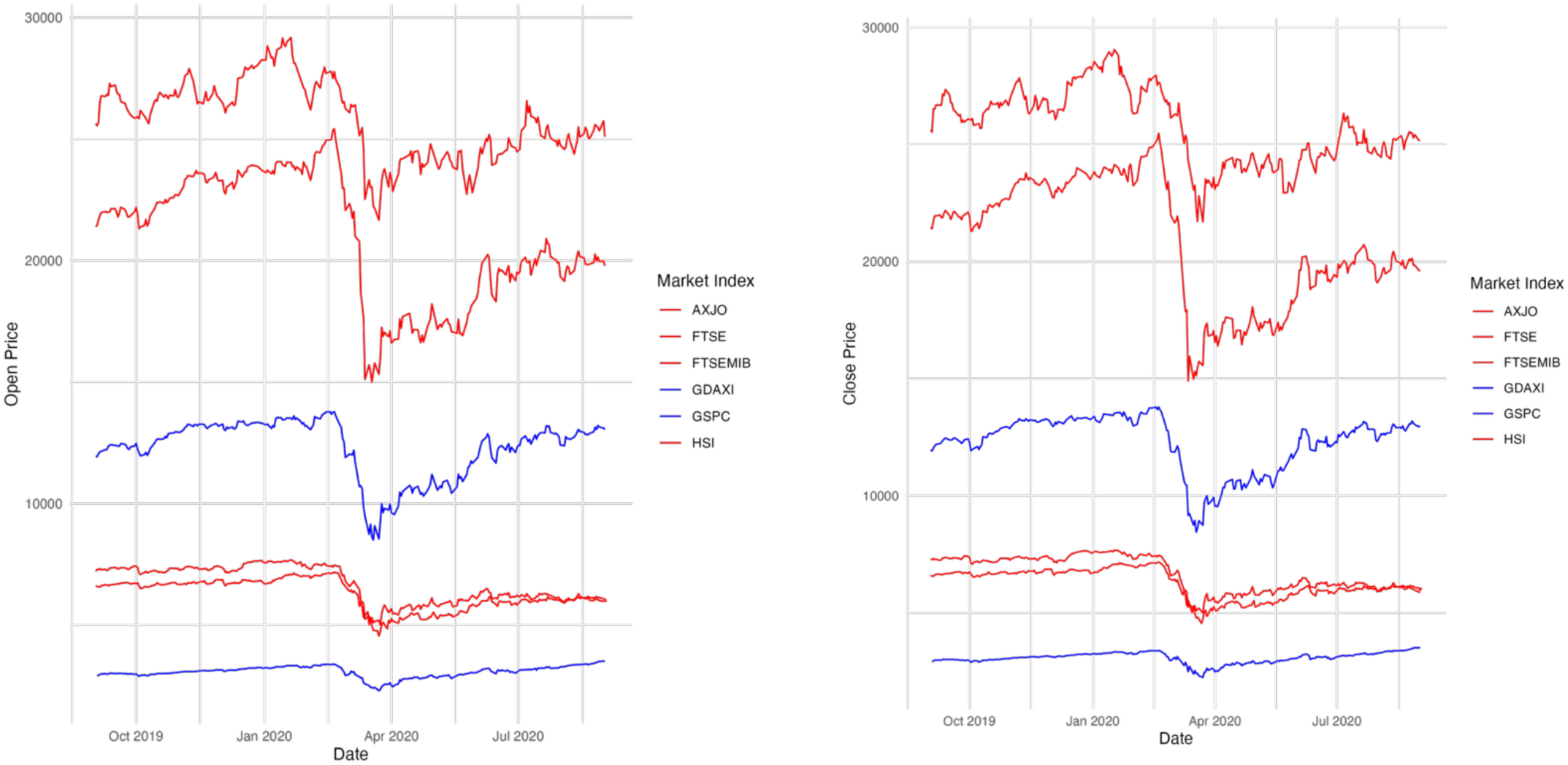
Shows the change in stock prices between September 2^nd^ 2019 and September 2^nd^ 2020, where red lines represent the stock prices that do not return to their original state after the onset of the pandemic and blue lines are stock prices that return or converge to prices prior to the onset of the pandemic. The left plot shows prices at open and the right plot prices at close. As will be demonstrated below, stocks that return to their pre-pandemic original prices in the sixth months following the onset of the pandemic, such as the GDAXI and the S&P 500, will not relate to the punctuated similarity models as the onset of dissimilarity will not have resulted in an effective change in stock prices.

## Experiment 2: Methods

### Data

For these models, stock price data was downloaded from the R ‘quantmod’ library (https://github.com/joshuaulrich/quantmod; last accessed: 17/06/2025). More specifically, this paper used downloaded stock price data between September 2^nd^ 2019 and September 2^nd^ 2020, therefore sampling price data approximately six months prior to and after the onset of the COVID-19 pandemic. To examine similarity in stock prices over this time period, the model was applied as in experiment one for the height data from children, where similarity was computed as 1-the difference between each pair of daily stock prices. This resulted in an average of 253 days from which to sample data (254 for the UK, 253 for the US, 252 for Germany, 253 for Italy, 256 for France, 255 for Australia and 249 for Hong Kong), therefore seven similarity matrices of ∼253 by ∼253 each. Stock price data is not recorded on Sundays and for some days data was missing, therefore, data does not exist for the exact number of days between September 2^nd^ 2019 to September 2^nd^ 2020. As in experiment 1, stock prices at open and stock prices at close were used as ‘replication’ and ‘discovery’ sets to confirm that modelled similarity was consistently related to stock price similarity. Thus, each market index resulted in two daily stock price similarity matrices, one based on the stock price at open and one based on the stock price at close (Figure 8).

**Figure 8:**
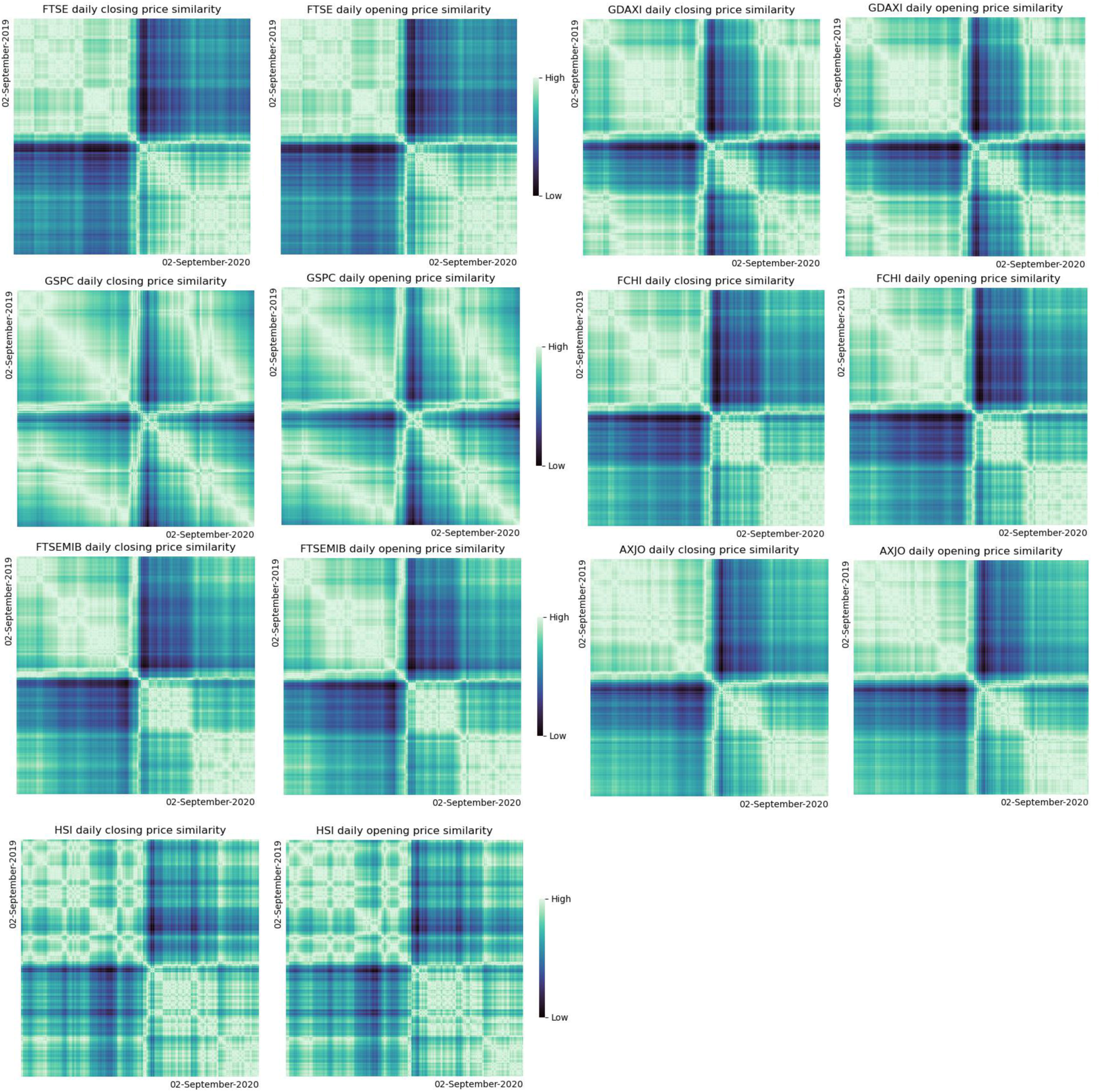
Real similarity distributions for stock prices, where each cell in the similarity matrix represents the similarity between the stock price at a given day to another and the diagonal represents perfect similarity (each stock with itself). Similarity plots on the left represent prices at closing and on the right prices at open. The similarity distributions are labelled by their related tickers. The FTSE-100 is labelled ‘FTSE’, the DAX as ‘GDAXI’, the S&P-500 as ‘GSPC’, the CAC 40 as ‘FCHI’, the FTSE-MBI as ‘FTSIMIB’, the S&P/ASX 200 as ‘AJXO’ and the HSI as ‘HSI’.

### Modelled similarity

Like in experiment 1, the hypothetical similarity models computed were nearest neighbor, convergence, divergence, punctuated similarity, and revised punctuated similarity models, which attenuated for nearest neighbor effects. To model similarity, the number of days was used from the start of the stock price data (time point 0 at the first recorded daily stock price), incrementing each day by the difference between dates at which stock prices were recorded. For example, if one were to start on the 2^nd^ of September 2020 and the next day recorded was the 3^rd^, the following on the 4^th^ etc., one would have time point 0 and then day 1, 2, 3 etc. But if there was a jump say from September 4^th^ to September 6^th^ (i.e., no stock price data was recorded on September 5^th^) then the days sequence would look like this: 0, 1, 2, 4 since September 6^th^ marks four whole days (not three) having passed since one started recording stock prices at time point 0.

Three key dates related to the COVID-19 pandemic as the central values for the punctuated similarity models were used. These were: the 13^th^ of January 2020, the day after China declared the existence of the SARS-CoV-2 virus (n.b. the actual date was the 12^th^ of January, which was a Sunday), the 11^th^ of March 2020, the day on which the World Health Organization declared COVID as a pandemic, and the 16^th^ of March 2020, the day on which the United States of America closed its border to non-US citizens or residents (Cucinotta & Vanelli, 2020; Spiteri et al., 2020). To note, other national lockdown dates were considered. However, to avoid the risk of over-testing very similar models, since national lockdown dates varied between the 9^th^ of March 2020 (Italy) and the 25^th^ of March 2020 (Germany), which were around the WHO and USA time points, time-points that may have had more consistent global impacts on stock market behavior were chosen. For each of the seven market indices examined, given that the time sequences were fairly equal, this paper only visualizes the modelled similarity values for times related to the French FCHI index, which had the highest number of daily stock prices recorded (N = 256) (Figure 9). The remaining plots for the other six indices can be found on the github repository for this project.

**Figure 9:**
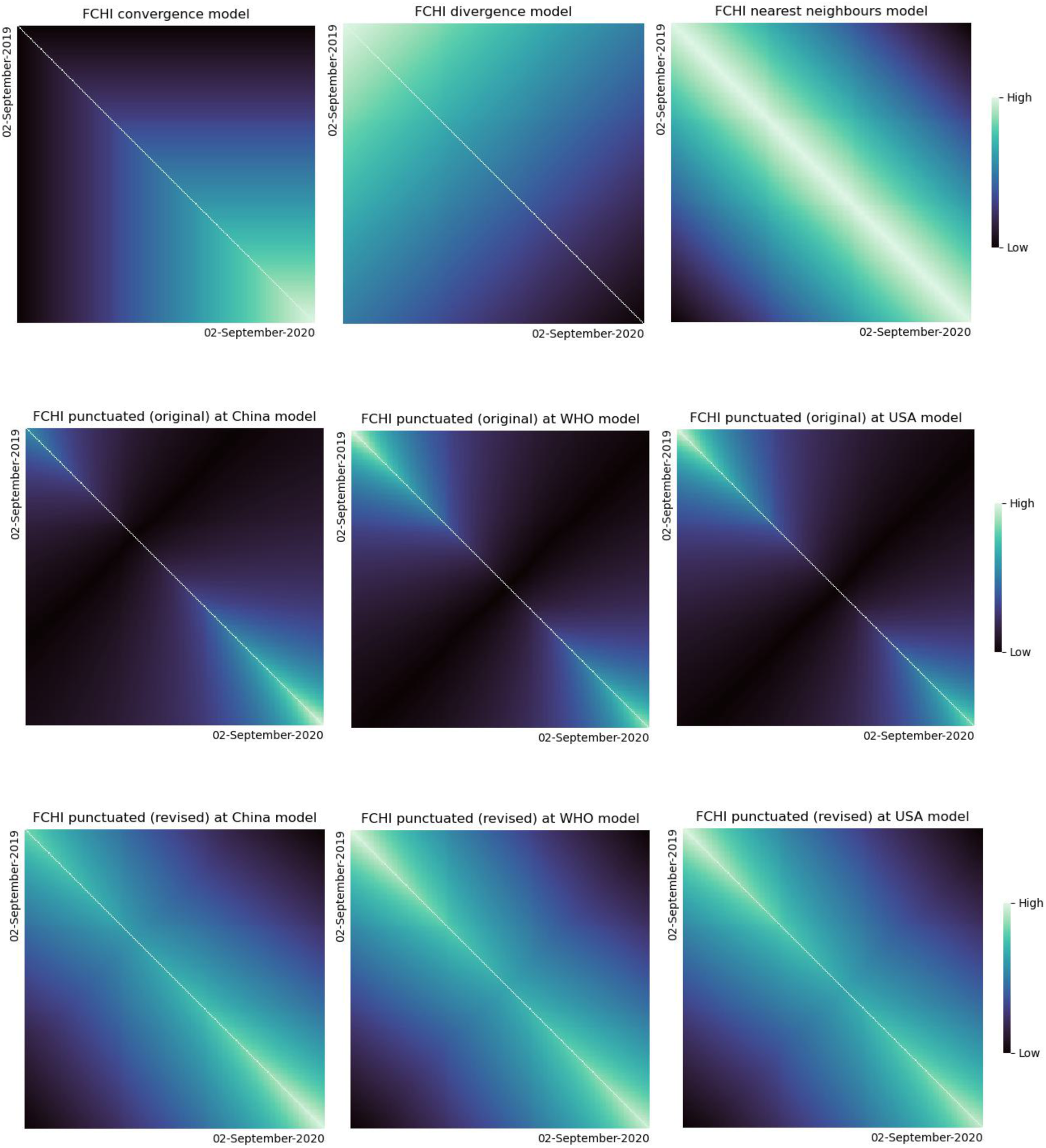
The modelled similarity plots for time between September 2019 and September 2020, measured as the increase in days from time point 0 (the upper left corner). In the upper row are the convergence, divergence and nearest neighbor models. The middle row presents the original punctuated similarity models and the bottom row the revised punctuated similarity models, attenuated for any nearest neighbor effects.

## Experiment 2: Results

Using the same methods as reported in experiment 1, results showed that for markets which did not immediately recover but were relatively stable in the 6 months after the onset of the COVID pandemic (UK, Italy, France, Australia, and Hong Kong), punctuated similarity was highly related to real similarity distributions for these markets (Table 3). Contrastingly, for markets (USA and Germany) that recovered in the sixth months after the onset of the pandemic and thus their stock prices returned to similar or higher prices as before the pandemic, price similarity did not relate to any of the modelled similarity distributions. Although the same follow-up analysis was run to determine which modelled similarity mostly related to the real distributions, the resulting R values were so low for the USA and Germany price similarity that nearly all models were returned as significantly high. Therefore, only the results from the mantel tests are reported (Table 4) to demonstrate that they never exceeded values of 0.21. For further inspection, raw outputs from this analysis are also available on the related git-hub repository.

**Table 3:**
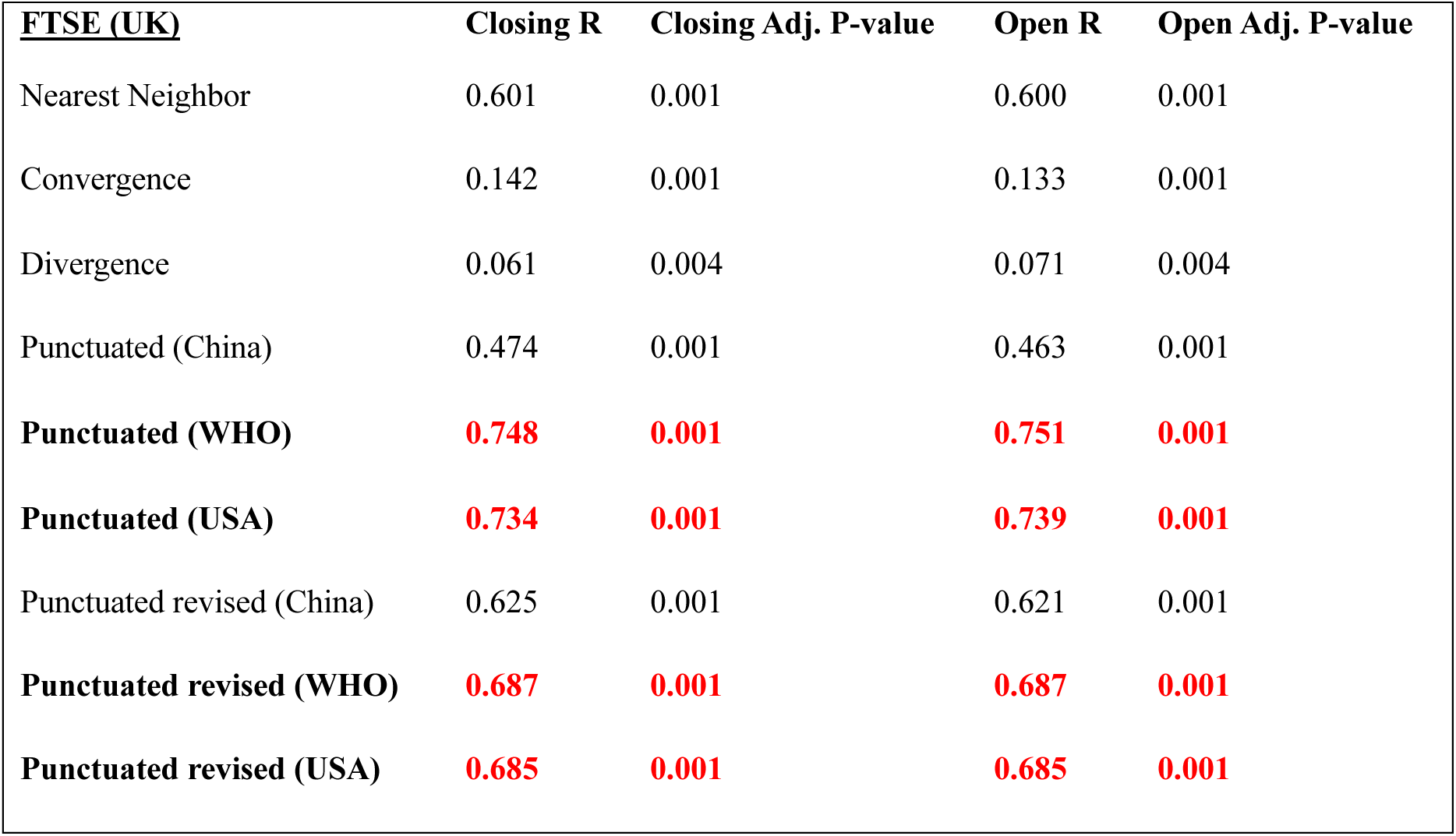
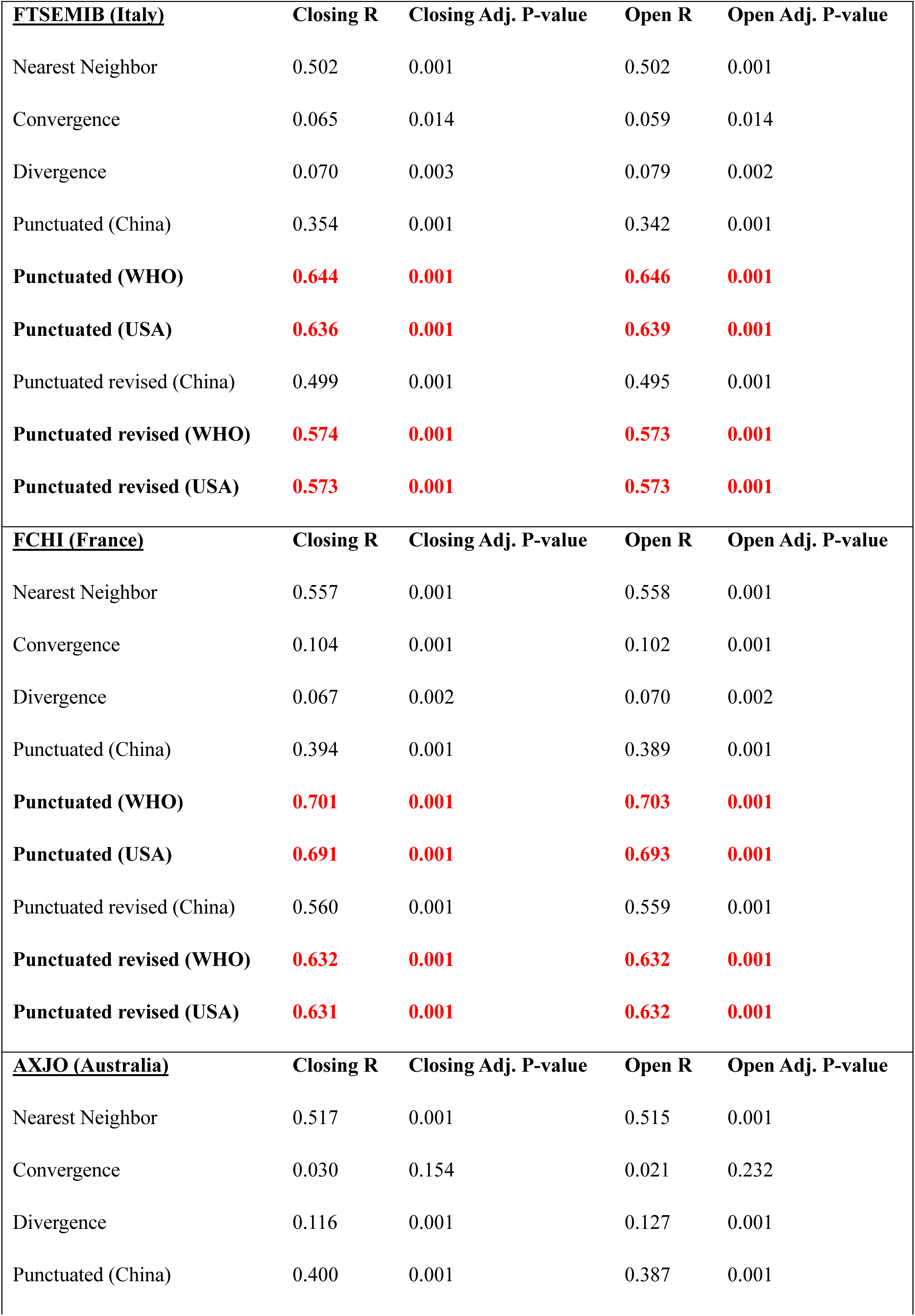
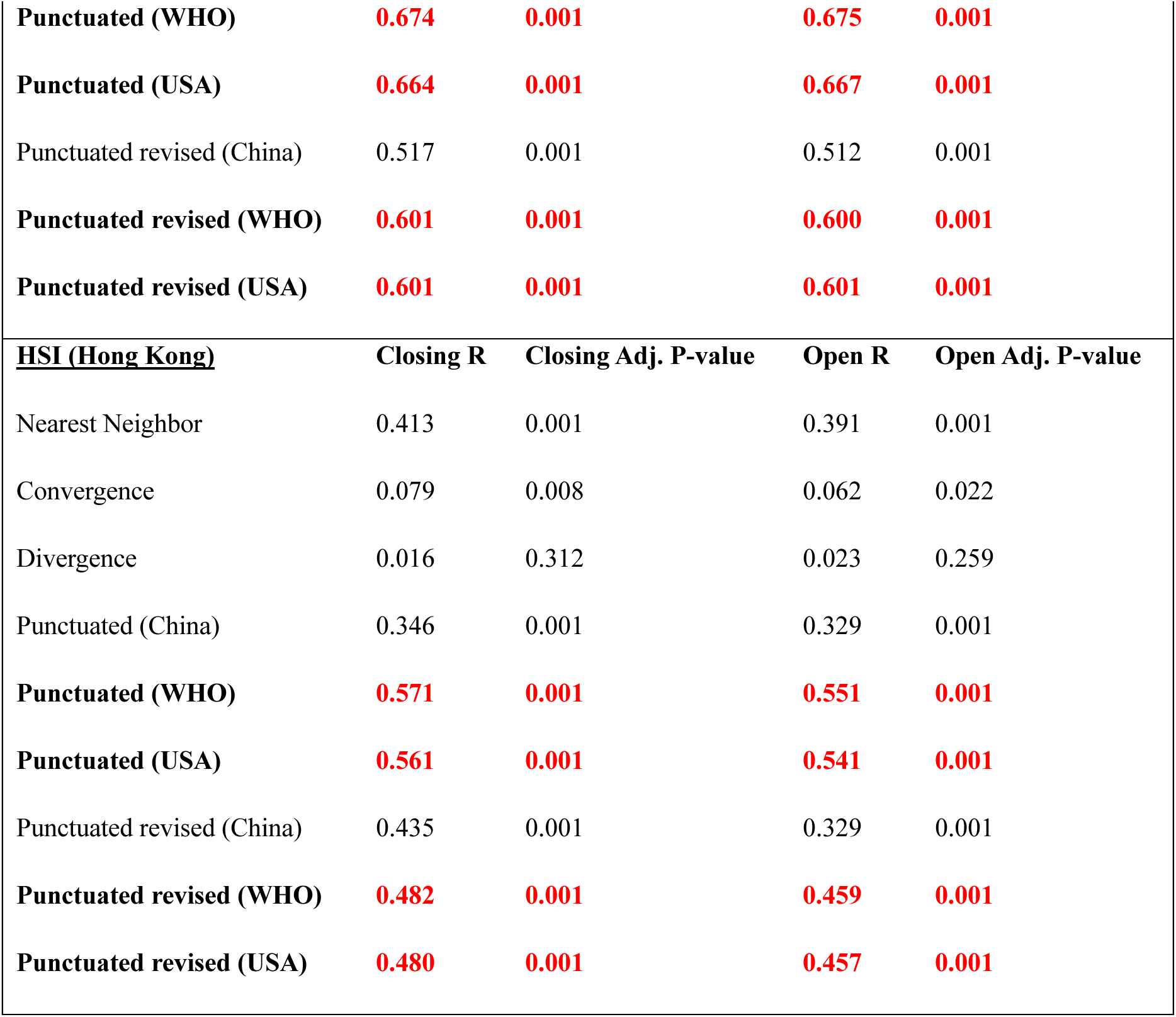
Results for markets that exhibited ‘punctuated’ behavior. *R values bolded in red were significantly higher than all other reported R values.*

**Table 4:**
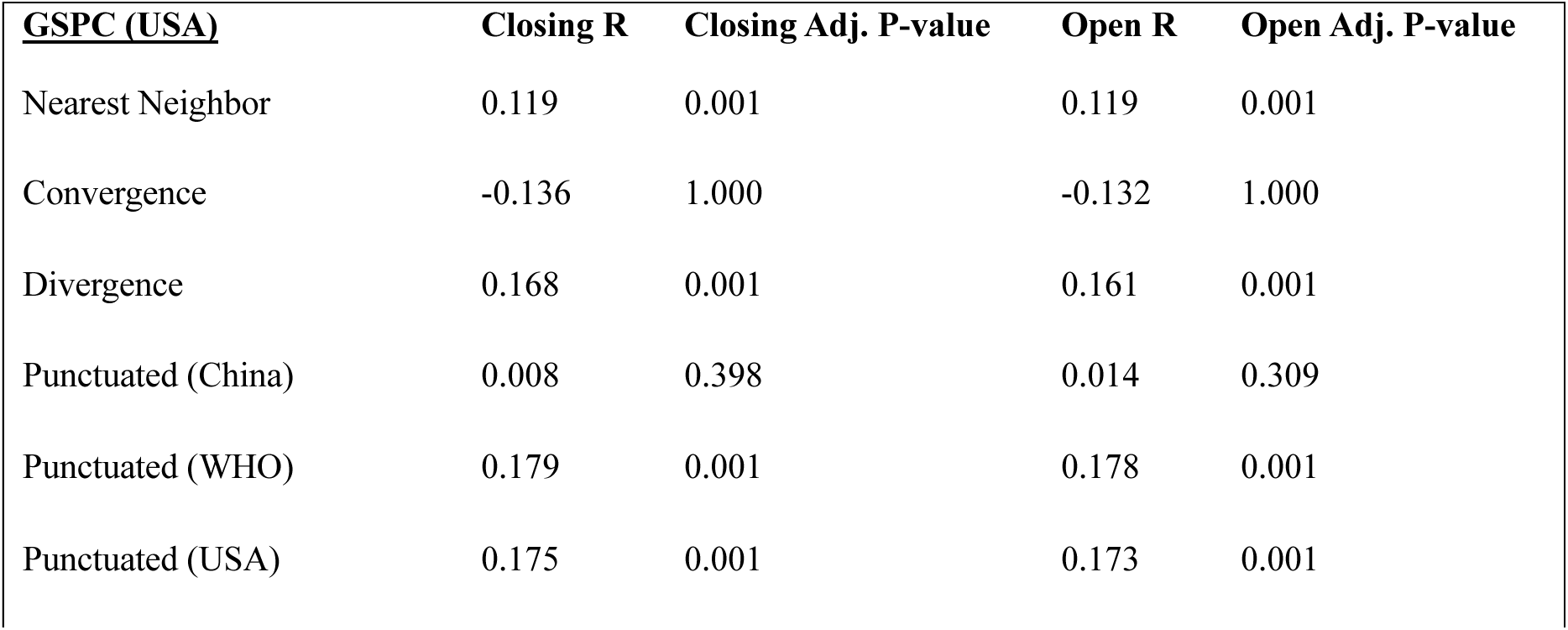
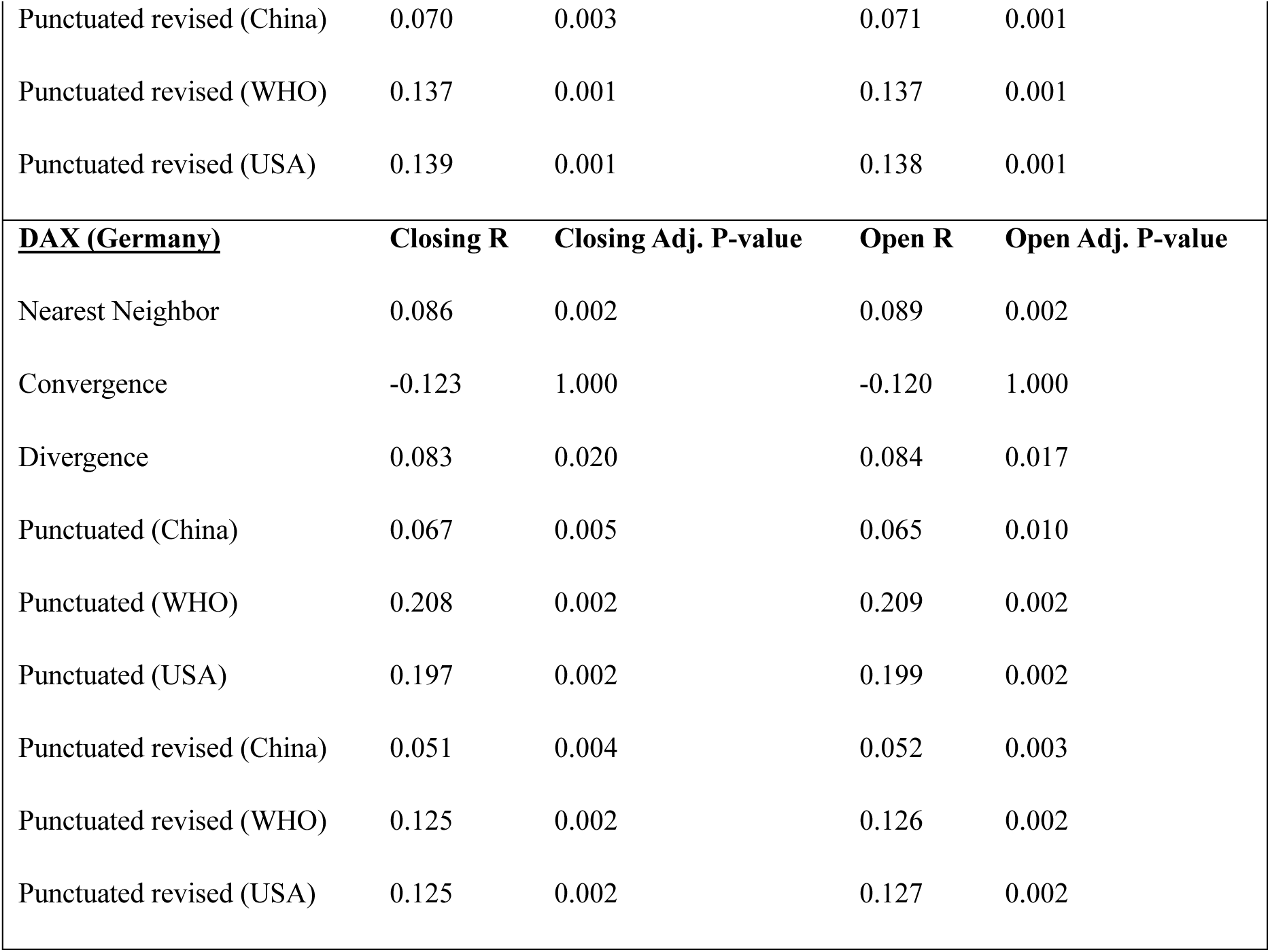
Results for markets that did not exhibit ‘punctuated’ behavior. *R values bolded in red were significantly higher than all other reported R values.*

## Discussion

This work presents a novel way to model similarity, where dissimilarity is modelled around a chosen value. Importantly, this model is flexible so that researchers can choose where they want to model dissimilarity, based on prior hypothetical assumptions about where dissimilarity might occur in their data, and whether they may want to consider possible nearest neighbor effects. Overall, this work demonstrates that the punctuated model is effective by showing that it expectedly relates to onset of height growth in puberty and to momentary stock market volatility. The model also strongly relates to prior notions of puberty being a pivotal moment for rapid changes in height, in line with previously reported differences in the onset of pubertal changes between boys and girls (Suutela et al., 2022), with boys relating to dissimilarity in height at later ages than girls. The punctuated similarity model also demonstrates that it is sensitive to critical moments related to stock price volatility (Kayani et al., 2024). Finally, this work is accompanied by an open-source R library to model the reported similarity distributions for future research usability.

Through stock market data, the punctuated model was further able to demonstrate that punctuated similarity is sensitive to data not returning to its original values after it has experienced a critical moment of dissimilarity. For example, while all children after puberty might be similar in height they are not similar again in height to children before puberty, as in, they do not return to their original state. In market dynamics, this would mean that where our punctuated similarity model applies, after stock prices have experienced a state of volatility, they may stabilize but do not return to or exceed their original stable price index before that moment of volatility. In sum, the punctuated similarity effectively captures ‘turning’ moments of dissimilarity that anticipate a stable change from one state to another. Being able to hypothetically confirm heightened variability in two unrelated data sets (childhood growth data and stock market prices) with similar properties (a known moment of increased variance in otherwise stable trends) suggests that this punctuated model has the potential to be applied to other data where one might expect such moments of nonlinearity.

One main limitation of these similarity models is that they do not provide insight into directionality. In the example of growth data, modelling similarity only tells us how height similarity changes over age. It does not tell us whether height increases or decreases as a function of age, so to know whether height increases with age we would need to first plot or empirically test that there is indeed an increasing relationship between height and age in childhood. In the example of stock market prices, modelling similarity does not tell us whether stock prices increase or decrease prior to or following a critical time-point of volatility. However, this limitation of similarity models is also their strength, where modelling similarity provides a readily explainable and complimentary approach to understanding complex, dynamic data (i.e., Atkinson et al., 2008; Butler & Shybalkina, 2025; Ponzi & Aizawa, 2000; Stasinopoulos & Rigby, 2008). In sum, the work proposed in this paper provides a methodological tool that, albeit being computationally taxing, can help researchers to confirm whether a prior expectation of heightened variance or volatility is present in their data at a known expected time point. Most importantly, the results obtained from the methods proposed in this paper are intuitive and easy to visualize, meaning they do not require a strong background in statistics or mathematics to be easily explained.

Another limitation to consider in this work is that mantel test p-values are unusually high, even after correcting for multiple comparisons. Therefore, this work’s approach of confirming statistically significant differences between resulting R values may be another way of controlling for whether relationships between modelled and real similarity are not only significant but highly significant against other such relationships. Thus, the resulting R values from the mantel test should be the clearest indicator of relationships between modelled and real similarity. For example, in the US and German markets, while R values were deemed significant by the mantel test it was evident that there were no strong relationships, as R values never exceeded 0.21. The mantel test was chosen in line with previous approaches (Finn et al., 2020; Camacho et al., 2023), however, going further it may be important to consider other analytical frameworks for comparing between two similarity distributions.

In conclusion, this work posits a novel way of approaching hypothesis testing in data, especially where one might lack experimental control over population or stimulus selection. Without such controls, data may be highly heterogenous or be characterized by brief moments of increased variance, such as in the growth data for children or the stock market data presented in this study. Hopefully, future research will be able to make use of the presented punctuated similarity models to further their own investigations. Importantly, this work hopes to have demonstrated a novel technique that leverages large-scale data heterogeneity while preserving traditional models of confirmatory, hypothesis testing practices in research in a flexible and dynamic way. The author welcomes comments and critiques from the wider research community as to how it may be possible to improve or further these similarity approaches.

## Data and Code availability

All the analysis in this paper can be reproduced following the code and data here: https://github.com/sarahkaarina/punctuated_similarity_proof. The analysis requires that the R library ‘similaritymodels’ be installed in the user’s R environment (https://github.com/sarahkaarina/similaritymodels).

## Acknowledgements

The work presented in this paper was completed while the main author was as a post-doctoral researcher at the Italian Institute of Technology. The work presented in this paper does not reflect the opinions of the Italian Institute of Technology. The author thanks Michael V. Lombardo for helpful discussions during the conceptualization of her ‘punctuated’ similarity models and Natasha Bertelsen for helpful feedback and comments on the manuscript draft.

